# Delivering mRNAs to mouse tissues using the SEND system

**DOI:** 10.1101/2023.01.28.522652

**Authors:** Masato Ohtsuka, Jurai Imafuku, Shuho Hori, Aki Kurosaki, Ayaka Nakamura, Tsubasa Nakahara, Takashi Yahata, Kolari Bhat, Steven T Papastefan, So Nakagawa, Rolen M Quadros, Hiromi Miura, Channabasavaiah B Gurumurthy

## Abstract

mRNAs produced in a cell are almost always translated within the same cell. Some mRNAs are transported to other cells of the organism through processes involving membrane nanotubes or extracellular vesicles. A recent report describes a surprising new phenomenon of encapsulating mRNAs inside virus-like particles (VLPs) to deliver them to other cells in a process that was named SEND (Selective Endogenous eNcapsidation for cellular Delivery). Although the seminal work demonstrates the SEND process in cultured cells, it is unknown whether this phenomenon occurs *in vivo*. Here, we demonstrate the SEND process in living organisms using specially designed genetically engineered mouse models. Our proof of principle study lays a foundation for the SEND-VLP system to potentially be used as a gene therapy tool to deliver therapeutically important mRNAs to tissues.

## Introduction

Nearly all mRNAs produced inside a cell are intended for translating proteins within the same cell. Recent findings show that some mRNAs produced in one cell are transferred to other cells and they express their protein products in the recipient cells. Intercellular mRNA transfer (IMT) is shown to have a special role in many biological processes, including embryological development and in neural signaling.^1,2^

IMT commonly occurs between adjacent cells via cytoplasmic exchange or membrane nanotubes ^3^. IMT can occur between cells located far away from each other via extracellular vesicles (EVs) ^4^. Two newer mechanisms of IMT were reported recently, involving encapsulation of mRNAs inside either proteins, ^5^ or inside virus-like particles (VLPs).^6^ The VLP-based RNA transfer phenomenon was named SEND (Selective Endogenous eNcapsidation for cellular Delivery).^6^ EV-based transport of RNAs has been known for nearly two decades and has received much attention from the scientific community; therefore, it is studied extensively. However, the concept of VLP-based RNA transfer is fairly new; at present there is only one research study on the SEND phenomenon.^6^

The SEND-VLP system involves three components: a PEG10 protein that has the property of binding to its own mRNA; a cargo mRNA (any protein-coding sequence flanked by untranslated sequences of Peg10 mRNA); and a fusogen such as vesicular stomatitis virus g protein (VSVg) or endogenous retroviral envelope-derived genes such as syncytin-A and syncytin-B. The PEG10 protein forms a viral-like coat around the cargo mRNA (because it will recognize the flanking untranslated region (UTR) sequences of Peg10 mRNA) to form VLPs. VLPs then exit the producer cell and attach to another cell (called the recipient cell) via a fusogen and deliver the mRNA to the recipient cell. Therefore, Peg10 can be engineered to package different protein-coding sequences (cargo) into VLPs, as long as they contain the flanking 5’ and 3’ UTR sequences of Peg10 mRNA.

In the pioneering work of Segel et al., 2021,^6^ the SEND system was demonstrated in cell culture model using Cre or Cas9 coding sequences as SEND cargoes. Plasmids of three SEND components (Peg10, cargo mRNA, and fusogen) were transfected to cells, and the VLPs produced from those cells were harvested and transduced to recipient cells to deliver the cargo. While this work elegantly demonstrates SEND in cultured cells, it is yet to be studied if SEND occurs in animals. We recently proposed several experimental strategies for testing SEND system in mice.^7^

## Results

In this work, we investigated whether the SEND-VLP process occurs in mice. We first constructed a plasmid-containing mRNA cargo expressing AkaLuc Luciferase. AkaLuc Luciferase is known to be one of the most sensitive bioluminescent receptors.^8^ The plasmid was administered to mice via a hydrodynamic delivery approach, along with the plasmids expressing the other two components of the SEND system; PEG10 and VSVg (fusogen). The mice were analyzed using in vivo imaging system (IVIS) two days after administering the SEND components (Fig. S1). We readily observed a bioluminescence imaging (BLI) signal in the liver (Table S1). The liver is one of the most common organs known to express proteins from plasmids delivered via the hydrodynamic method.^9^ The BLI signal in liver cells could be from one of two origins: from the expression resulting directly from the cargo plasmid or from a cargo mRNA version produced as encapsulated VLPs, which are delivered to (and expressed in) cells (referred as VLP recipient cells). Since we did not observe a BLI signal in any tissues other than liver, it is hard to conclude whether the BLI signal in (all of) the liver cells was from the plasmid source alone or if it was also from VLP-mediated IMT in some cells.

The reasons we could not detect a BLI signal in tissues other than liver (which is indicative of IMT) may be due to some limitations in these experiments: (a) the amount of exogenously supplied plasmids (of three different components) may not be adequate to elicit a detectable level of VLP-mediated IMT; (b) the sensitivity of the system may not be high enough to detect cells that may have received mRNA cargo via IMT in tissues other than liver, especially if such cells are very few; (c) the timing of analyzing mouse tissues following hydrodynamic delivery and after substrate administration, including the dose of substrate, may need to be optimized to obtain detectable levels of VLP-mediated IMT; and lastly, (d) as stated above, it is difficult to conclude whether some of the BLI signal in liver was a result of the mRNA cargo of VLPs received by cells or if it was solely because of direct expression from the cargo plasmid.

We then set out to test the SEND system by delivering Cas9 mRNA as a cargo, with a readout of gene editing to correct a frameshift-edited GFP (ΔEGFP) locus in a CRISPR-Cas reporter mouse model we recently developed.^10^ Testing the SEND system using the ΔEGFP reporter model requires, apart from the SEND components, the administration of one additional reagent: a guide RNA targeting the frameshifted GFP sequence (EGFP-Cr1). We administered plasmids expressing Peg10, VSVg, Mm.Cargo (SpCas9), and synthetic two-part guide RNA (crRNA and tracrRNA) to two ΔEGFP mice. In both mice, we observed very few cells showing GFP fluorescence in the liver (Table S2). This result was expected, considering that plasmids and guide RNA components may not get delivered to the same set of cells, which is ultimately needed for successful editing of frameshifted GFP to restore its fluorescence.

To overcome this limitation, and to ensure a uniform supply of guide RNA in all cells of the mouse, we generated a knock-in mouse expressing single-guide RNA (sgRNA), targeting the frameshift mutant region of the ΔEGFP mouse (EGFP-Cr1)^10^ (Fig. S2). The ΔEGFP/Cr1 model was developed using the CRISPR-based methods and guide RNAs previously described.^11–16^ The two mouse models (ΔEGFP and EGFP-Cr1) were bred together to generate a double mutant strain (ΔEGFP/Cr1). The above experiment (delivering Cas9 mRNA using SEND) was repeated on the ΔEGFP/Cr1 model by administering only the SEND component plasmids (PEG10, VSVg, and Mm.Cargo[SpCas9]). A schematic of the ΔEGFP/Cr1 mouse model and the expected outcomes of hydrodynamic delivery of SEND is shown in Figure 1A. Mice were administered the above plasmids via hydrodynamic delivery and analyzed at seven days post delivery (Fig. 1B). Representative images of one of the two mice analyzed are shown in Figures 1C and 1D. The results showed fluorescence in many of the liver cells (Fig. 1C). Of note, we also detected fluorescence in spleen cells in both mice (Figs 1D and S3, supplementary text 1, supplementary movie M1 and Table S3). We repeated the experiment on three more mice, and analyzed them at days 8, 13 and 60 day post hydrodynamic delivery (Fig S3). All animals showed reproducible results; fluorescence in many liver cells and spleen cells. From these experiments, we conclude that the fluorescence in spleen cells could very likely have originated from the VLPs produced in liver traversed to spleen cells resulting in IMT.

**Figure 1.**
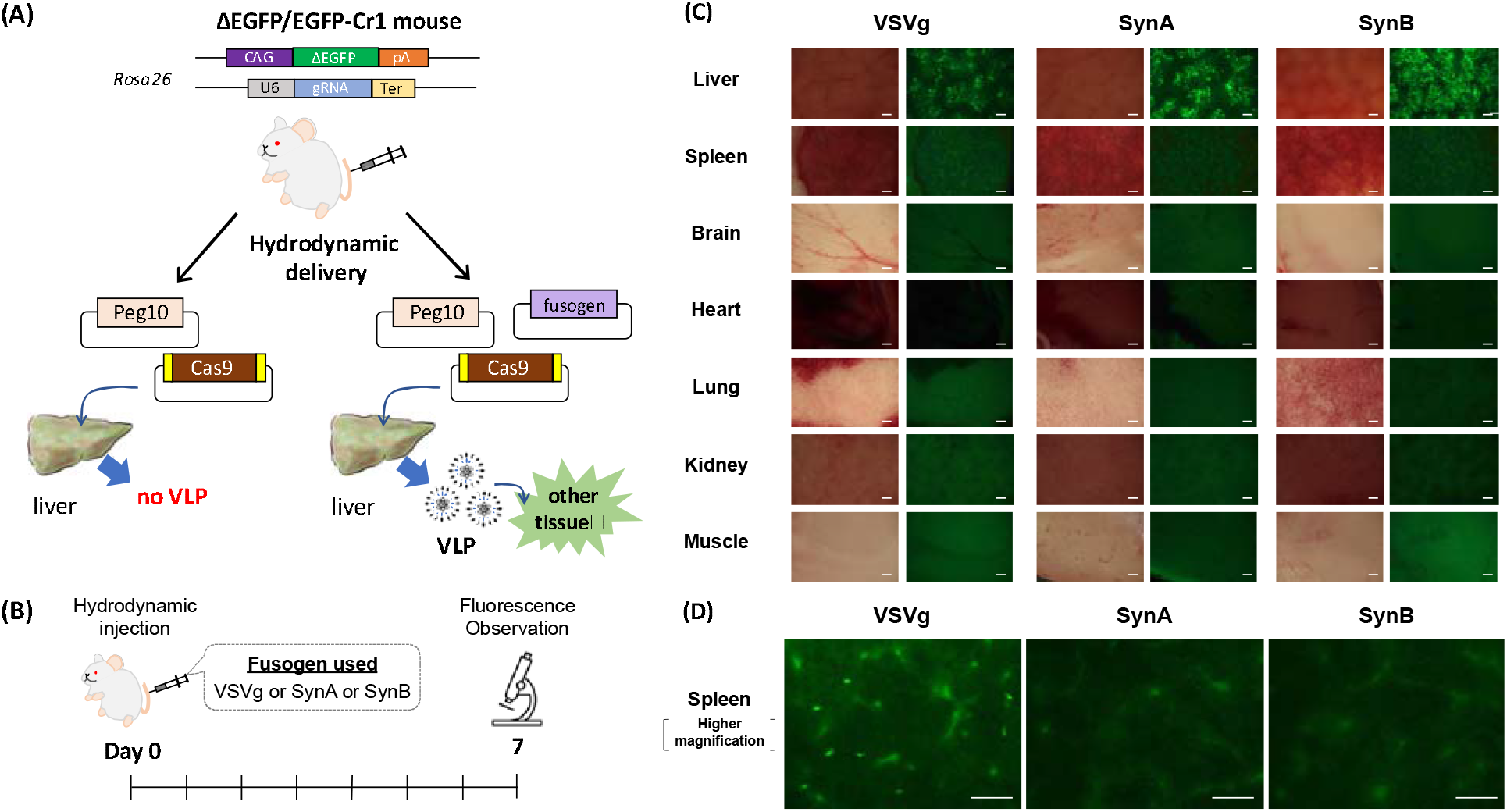
Validation of SEND system using genetically engineered mouse models. (A) schematic of trans-heterozygote mouse containing one allele each of ΔEGFP reporter and EGFP-Cr1 gRNA, at the ROSA26 locus. The two alleles provide the genetic components needed for functional readout for the SEND system, delivering an mRNA cargo coding for CAS9 protein. Three SEND plasmids to be delivered code for Cas9 mRNA cargo, PEG10, and fusogen proteins. GFP fluorescence in liver indicates the expression of CAS9 protein, leading to restoration of frameshifted EGFP. It is well known that the liver cells typically express genetic components directly from plasmids; therefore, it is hard to differentiate if some of the fluorescent liver cells were because of the SEND-VLP mode of IMT. Because plasmids delivered from tail vein via hydrodynamic mode do not generally express in other tissues, the presence of fluorescence in tissues other than liver likely indicates SEND-VLP mode of IMT. (B) Hydrodynamic treatment regimen. (C) Fluorescent stereomicroscopy of tissues from a representative mouse analyzed at seven days (time point). (D) Fluorescent microscopy images of spleen tissue showing punctate fluorescence. Three fusogens were tested—VSVg, syncytin-A (SynA), and syncytin-B (SynB), of which VSVg showed fluorescence in spleen. The fluorescence in spleen is likely due to SEND-VLP mode of IMT (see text for discussion, supplemental movie S1 and supplemental text 1). Bar = 200µm.

## Discussion

Our conclusion that the fluorescence observed in spleen cells could be a result of IMT is based on the following additional points. We could detect fluorescence only when the sensitivity of the experimental system was addressed by using a genetic model especially suitable for testing the SEND system; the model that uniformly expresses ΔEGFP reporter and sgRNA in all tissues. However, none of the other experiments showed fluorescence in tissues other than liver, even when the cargo plasmid concentration remained constant in all experiments. In other words, if the cargo expression in spleen cells originated directly from the cargo plasmid (instead of VLP), fluorescence in spleen cells should have been noticed in other experiments too; for example, those experiments involving delivery of exogenous guide RNA to ΔEGFP mouse and those involving other fusogens (see below). To verify whether our conclusion was correct, we delivered only the cargo plasmid Mm.cargo(SpCas9) and excluded Peg10 and VSVg fusogen plasmids from the mixture of hydrodynamic delivery components (Table S4). As expected, we observed fluorescence only in liver; in this experiment, other tissues did not show fluorescence possibly because VLPs would not be generated without exogenously supplied PEG10 and a fusogen. We repeated the experiment by changing the fusogen from VSVg to SynA or SynB. We did not detect fluorescence in spleen with either of those fusogens (Figs. 1C and 1D). Of note, VSVg is shown to have highest effect in a cell culture assay system, whereas SynA showed a weaker effect than VSVg, and SynB did not show any effect in the initial report by Segel et al. in 2021.^6^

We do not know if the sensitivity of the assay (using SynA or SynB fusogens) was sufficient enough to detect fluorescence or whether these fusogens are not involved in the specific experimental context. We speculate that the reason could be because of the lack of sensitivity of our assay (rather than the non-involvement of fusogens). This is based on the following experiments and observations reported by others. Prior reports demonstrate that SynA and SynB-pseudotyped lentiviral vectors could be transfected into primary mouse splenocytes with relatively high efficiencies whereas lentiviral delivery to murine spleen in vivo was only moderately efficient ^17^. The absence of transduction in spleen cells in our experiments with SynA or SynB was possibly because the effect of VLPs (unlike lentiviral vectors) was too weak for detection. Related to this, Segel et al. 2021 observed reduced efficiency of VLPs in comparison to lentiviral vectors for Cas9 mRNA delivery, as well as reduced efficiency of syncytin-pseudotyped versus VSVg-pseudotyped VLPs. Thus, we presume that the combination of these two factors (i.e., lower efficiency of VLPs vs lentiviral vectors and the lower efficiency of SynA or SynB fusogens compared to VSVg fusogen) may have resulted in low efficiency transduction that was below the detection threshold of our experimental methods. Further, the receptor for SynA, lymphocyte antigen 6 family member E (Ly6e), is highly expressed in murine spleen (in addition to liver, lung, and placenta) and mediates cell-cell fusion upon binding with SynA,^18–20^ which suggests that fusogens like SynA and SynB could be involved in SEND-VLP phenomenon in spleen but again our experiments may not have been sensitive enough to detect it. In addition, recent studies have reported several genes that reduce cell-cell fusion efficiency of endogenous retroviral envelope genes by interfering their receptors in humans ^21,22^ and various primates.^23^ While no interfering elements have been reported for SynA and SynB, they may also be involved in transduction efficiency. Further studies using in vitro and in vivo models would be helpful in understanding the role (or involvement) of SynA or SynB fusogens in SEND based VLP biogenesis. Along those lines, experiments involving direct injection of exogenously produced VLPs containing fusogens SynA or SynB to mice may achieve a higher sensitivity (than the plasmid administration experiments) for detecting cargo delivery to spleen cells.

In summary, the two sets of experiments involving no fusogen (Table S4) and weaker or ineffective fusogens (Figs. 1C and 1D and S3) further strengthens our conclusion that the fluorescence in spleen we observed (with the use of VSVg fusogen; Fig. 1D) was likely a result of VLPs formed in liver that traversed to spleen.^24^ It is known that hydrodynamic delivery elicits about 10,000-fold higher expression of plasmids in liver cells compared to cells in other tissues ^25,26^.

While our experiments are not quantitative, the observation that a few cells in spleen show fluorescence indicate the strong possibility of the SEND-VLP mode of IMT. Even though we devised a sensitive approach to demonstrate the SEND system in a mouse genetic model, one of the limitations of relying on gene-edited restoration of GFP as the readout of our experiment is that only a third of the gene edits can elicit fluorescence because edits that result in two of the three reading frames remain undetected. Some of the reading-frame restored edits would not result in fluorescence because the amino acid change may not produce fully functional GFP. Of note, in our previous study (using the ΔEGFP reporter mouse model),^10^ we observed that only 20% of all indels were reading-frame restored gene edits.

Not much is known about the SEND-VLP system. The only published report demonstrates the process using in vitro cultured cells. Also, there are no tools available to study the SEND phenomenon in an in vivo setting. mRNA-based therapeutics and vaccines, and LNP-based tools to deliver those, have been gaining popularity and are already being successfully used in humans.^27^ The SEND system has broad potential for clinical application given the ability to guide tissue tropism based on alternative pseudotyping strategies and the potential for reduced immunogenicity given its endogenous components.^28^ For example, in utero gene therapy, which involves therapeutic mRNA delivery to the fetus or placenta to ameliorate developmental anomalies, would greatly benefit from mRNA delivery strategies involving entirely endogenous components that target the placenta, such as with syncytin fusogens.^29^ Additionally, further study of the mechanisms of SEND-based mRNA delivery may lead to an improved understanding of cell-cell communication in development and between various organ systems. While our proof of principle study lays a foundation that SEND-VLP system can be used as a gene therapy tool for delivery of mRNAs expressing therapeutically important proteins to tissues, the detailed cellular mechanisms of SEND can be understood using genetically engineered mouse models containing individual components of the SEND system.^7^

## Materials and Methods

### Plasmid constructs

The following Addgene plasmids were used in this study: pCMV-MmPeg10rc4 (174858), pCMV-SynA (174860), and pCMV-Mm.cargo(SpCas9) (174865). The pCMV-Mm.Cargo(Akaluc) plasmid was constructed by inserting a gBlock sequence coding for Akaluc via Gibson assembly into the AgeI and BsrGI enzyme sites of the Addgene plasmid 174862 (pCMV-Cargo(CRE). This cloning step replaces the parent plasmid’s Cre coding sequence (1047 to 2093 bases) with the Akaluc coding sequence. The construct was generated using the cloning services from Genepass, Tennessee. Complete sequence of the final construct was confirmed with services from Plasmidsaurus, Eugene, OR. The pBNA-(CMV-VSVg) plasmid was generated by excising out ‘Rev’ region of the pCMV-VSV-G-RSV-Rev (RDB04393) using the RE enzyme Pvull. All plasmids were purified using the Endotoxin-Free Plasmid Purification Kit (QIAGEN) prior to the *in vivo* gene-delivery experiment.

#### A note about the pCMV-MmPeg10cr4 plasmid

When we compared the mPeg10 amino acid sequence of the Addgene plasmid 174858 with that on the mouse genome (NP_570947.2), all amino acids were identical excluding two deletions (positions at 685-690 and 698-745) and nine amino acids insertion of Hemagglutinin (HA) tag at the C terminal (**Fig. S4**). The two deleted regions were found to be part of a disorder region predicted by MobiDB-lite^30^ and they were not part of any protein motifs of retroelements as analyzed using the InterPro^31^ and the Gypsy Database 2.0^32^ searches. Considering that the HA tag at the C terminus of PEG10 did not affect its function in a previous study^6^ and considering our analysis that the two short deletions were not part of any protein motifs, we conclude that the Addgene MmPeg10 construct (174858) used in this study may not affect the function of the protein, and thus allow for intercellular mRNA transfer (IMT) when used with VSVg.

### Generation of ΔEGFP/Cr1 mouse

#### EGFP-Cr1 mouse

The EGFP-Cr1 mouse was a knock-in mouse model, at the ROSA26 locus, carrying an expression cassette of guide RNA against a frameshifted region of a ΔEGFP mouse. The model was generated using the *i-*GONAD method.^11,12^ Briefly, a solution containing tracrRNA, crRNA, and Cas9 protein, as well as long single-stranded DNA (ssDNA) as a donor repair template ^13,14^, was injected into the oviduct of 0.7 day old pregnant MCH (ICR) strain female followed by in situ electroporation. The 500 bases long ssDNA donor (gift from GenScript) contained the U6 promoter-driven EGFP-Cr1 sgRNA expression cassette and the ROSA26 homology arms. The guide RNA used for inserting the cassette was ROSA-Cr4, which was described previously ^16^. Offspring with knock-in mutations were identified using PCR ^15^ and sequencing-based genotyping strategies described previously ^13,14^.

#### Modifying the coat color of the ΔEGFP mouse strain

Compared to a black coat color, an albino coat color is better for easily finding the tail vein needed for intravenous administration of reagents. Therefore, we changed the coat color of our recently developed ΔEGFP reporter mouse ^10^ from black to albino as follows. First, we edited the tyrosinase gene of the C57BL6 strain using a CRISPR guide AACTTCATGGGTTTCAACTG [CGG (PAM)]. The *i-*GONAD ^11,12^ method was used for creating the mutation. The injection solution contained tracrRNA, crRNA, and Cas9 protein. The offspring were assessed by albino coat color, indicative of successful gene editing. Mice were also genotyped by PCR and sequencing of the target site. An albino offspring with three base deletion was bred with the ΔEGFP mouse to obtain the albino ΔEGFP mouse strain.

The EGFP-Cr1 and the albino ΔEGFP mice were bred together to obtain the albino ΔEGFP/Cr1 strain that was used for the SEND experiments.

#### Hydrodynamic delivery

Hydrodynamic delivery was performed as described previously ^25^. Briefly, animals were placed in the mouse holder (Braintree Scientific, Inc.) and were injected with a solution (one-tenth of the mouse weight in volume; for example, 2.3 ml/23 g mouse) containing SEND plasmids (1.1 μg/ml for each plasmid) using a 3 ml luer-lok type syringe (Becton Dickinson and Company [BD]) fitted with a 27-gauge needle (BD). Injections were performed at a constant speed via tail vein and completed at around five seconds. In the hydrodynamic delivery experiment of CAS9 protein (Fig. S2C), CAS9 protein solution was injected without gRNA.

#### In vivo bioluminescence imaging

In vivo bioluminescence imaging (BLI) was performed using the IVIS Lumina II (PerkinElmer). Mice treated with hydrodynamic injection were anesthetized using 2.5% isofluorane, and 5 mM Akalumine-HCl (Sigma-Aldrich) substrate was injected intraperitoneally. The images were acquired 4 min (*in vivo*) and 17 min (*ex vivo*) after the substrate administration. The following conditions were used for image acquisition: an open emission filter, exposure time of 30s (*in vivo*) or 60s (*ex vivo*), binning 2, field of view 24 and f/stop 2. Data was analyzed by Living Image 4.3 software (PerkinElmer) specialized for IVIS system.

#### Fluorescence observation

Tissues were collected for microscopy after euthanizing the animals. Animals were euthanized as per the institute approved protocol. The EGFP fluorescence in various tissues, including liver and spleen, was observed under fluorescence stereomicroscope, with a GFP filter, and the fluorescent images were acquired using EOS Kiss X10. In some experiments (e.g., Fig 1C), spleen tissues were observed under inverted fluorescence microscope, with U-MNIBA and U-MWU filter cubes, and the fluorescent images were acquired using EOS Kiss X5.

## SUPPLEMENTARY MATERIALS

Figs. S1 to S4

Tables S1 to S4

Supplementary text 1

Movies S1-S3.

### Supplementary figures

**Figure. S1**. Hydrodynamic delivery of SEND components containing Akaluc mRNA cargo. (A) Schematic of the Mm.Cargo(Akaluc) plasmid. (B) Regimen of hydrodynamic treatment- and analysis-of mice (C) Representative IVIS images of mice analyzed. Internal organ (top to bottom): Heart and Lung, Liver, Spleen, Kidney, Intestine, and Muscle.

**Figure S2**. Development of EGFP-Cr1 mouse. (A) Schematic of the *i-*GONAD based method to insert the guide RNA expression cassette at the mouse ROSA26 locus. (B) Sanger sequencing electropherogram showing correct insertion of the donor ssDNA cassette. (C) Representative fluorescent microscopy of liver tissues after injection of CAS9 protein only (+Cas9: without gRNA, -Cas9: no injection control). Bar = 200µm.

**Figure S3**. Additional SEND experiments using the ΔEGFP/Cr1 mouse. (A) Hydrodynamic treatment regimen and the number of mice analyzed at each time point. (B) Fluorescent microscopy of various mouse tissues analyzed at various time points (see text for discussion). Bar = 200µm.

**Figure S4**. Pairwise alignment of mPeg10 amino acid sequences of the Addgene plasmid 174858 and that on the mouse genome (NP_570947.2). A retrotransposon gag protein motif (Pfam domain: PF03732) or a consensus disorder predicted region predicted by MobiDB-lite were indicated by red or blue line, respectively. The two deletions and an HA tag insertion in the Addgene plasmids are shown with red and blue open boxes respectively.

### Supplementary text 1

As part of screening GFP fluorescence (which is indicative of SEND-mediated IMT), we imaged the whole organs/tissues of various mouse tissues. However, background fluorescence is one of the biggest challenges when whole organs are imaged. Therefore, we do not know if we missed true GFP fluorescence in some of the tissues. However, we noticed clear punctate GFP fluorescence in at least spleens in all the five mice that were subjected to VSVg fusogen-mediated SEND experiments. Even though the still images (shown in Figure 1D) indicate GFP positive cells in spleen, background can still obscure true signal. In order to rule out background signal, we further analyzed the still images that were imaged with U-MNIBA filter cube or with U-MWU filter cube by creating movies using the iMovie tool. Three movies, S1, S2 and S3 (for VSVg-or SynA-or SynB-fusogens respectively), are included as supplementary movies. Each movie constitutes two representative regions of spleens captured with U-MNIBA filter (for GFP) or U-MWU filter (for negative control). Note that, of the three fusogens, only VSVg resulted in SEND-mediated IMT in spleen and thus the still images and movies of SynA and SynB fusogens serve as negative controls. The movies help in distinguishing background versus true signal more clearly compared to still images.

**Movie S1:** Movie of still images of two representative regions of tissues imaged with U-MNIBA (for GFP) or U-MWU (for negative control) filters, for the experiment with fusogen VSVg.

**Movie S2**: Movie of still images of two representative regions of tissues imaged with U-MNIBA (for GFP) or U-MWU (for negative control) filters, for the experiment with fusogen SynA.

**Movie S3:** Movie of still images of two representative regions of tissues imaged with U-MNIBA (for GFP) or U-MWU (for negative control) filters, for the experiment with fusogen SynB.

## Acknowledgments

pCMV-VSV-G-RSV-Rev plasmid (used for generating the VSVg fusogen construct) was procured from RIKEN; we thank late Dr. Hiroyuki Miyoshi for developing the plasmid and depositing the plasmid to RIKEN. We thank Joe Miano, Augusta University, USA for critical reading of the manuscript. We thank Nick May, TypeRight, for copy editing. CBG is funded by NIH grants R35HG010719, R21GM129559, R21AI143394, and R21DA046831. MO is funded by JSPS KAKENHI (20K21551) and Takeda Science Foundation (2020).

## Author contributions

MO and CBG conceived the overall idea, designed experiments, supervised the study, analyzed results, and wrote the manuscript. MO and HM designed and generated EGFP-Cr1 mouse strain. HM, STP and SN wrote some parts of the methods and discussions sections. JI performed hydrodynamic injections. MO, JI and SH collected microscopy data. SH collected IVIS data. AN performed i-GONAD experiments and TN performed some mouse breeding experiments. MO, HM and SH compiled and analyzed microscopy data (fluorescence and bioluminescence). AK, TY, KB and RQ generated plasmid constructs.

## Competing interests

A patent application based on this work is under preparation in which MO and CBG will be listed as inventors.

