## Supplementary material for "Delivering mRNAs to mouse tissues using the SEND system": Fig S1

### Slide 1
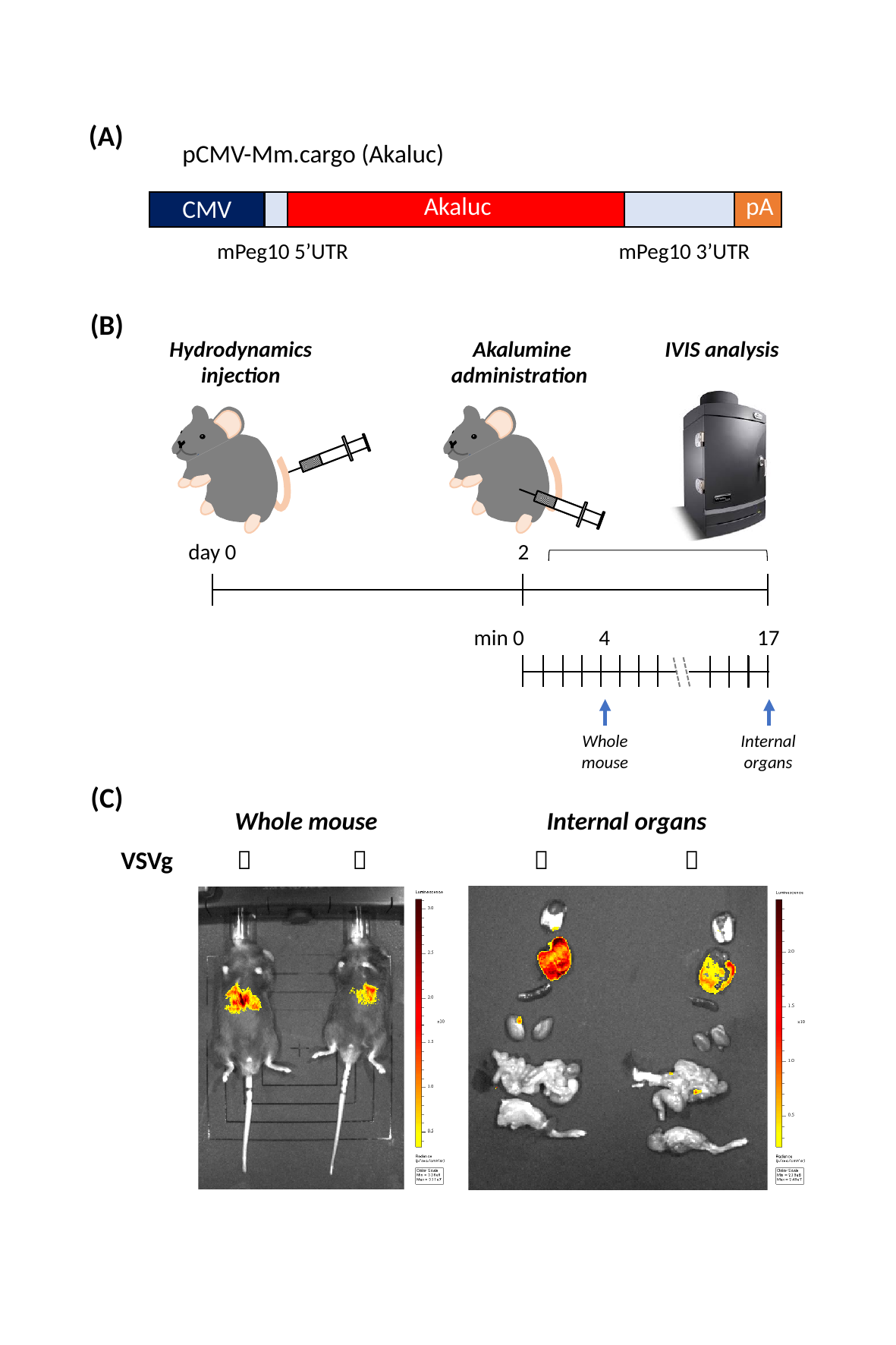

(A)
pCMV-Mm.cargo (Akaluc)
pA
Akaluc
CMV
mPeg10 5’UTR
mPeg10 3’UTR
(B)
Hydrodynamics
injection
Akalumine
administration
IVIS analysis
day 0
2
min 0
4
17
Whole
mouse
Internal
organs
(C)
Whole mouse
Internal organs
VSVg
＋
ー
＋
ー
