## Supplementary material for "Delivering mRNAs to mouse tissues using the SEND system": Fig. S2

### Slide 1
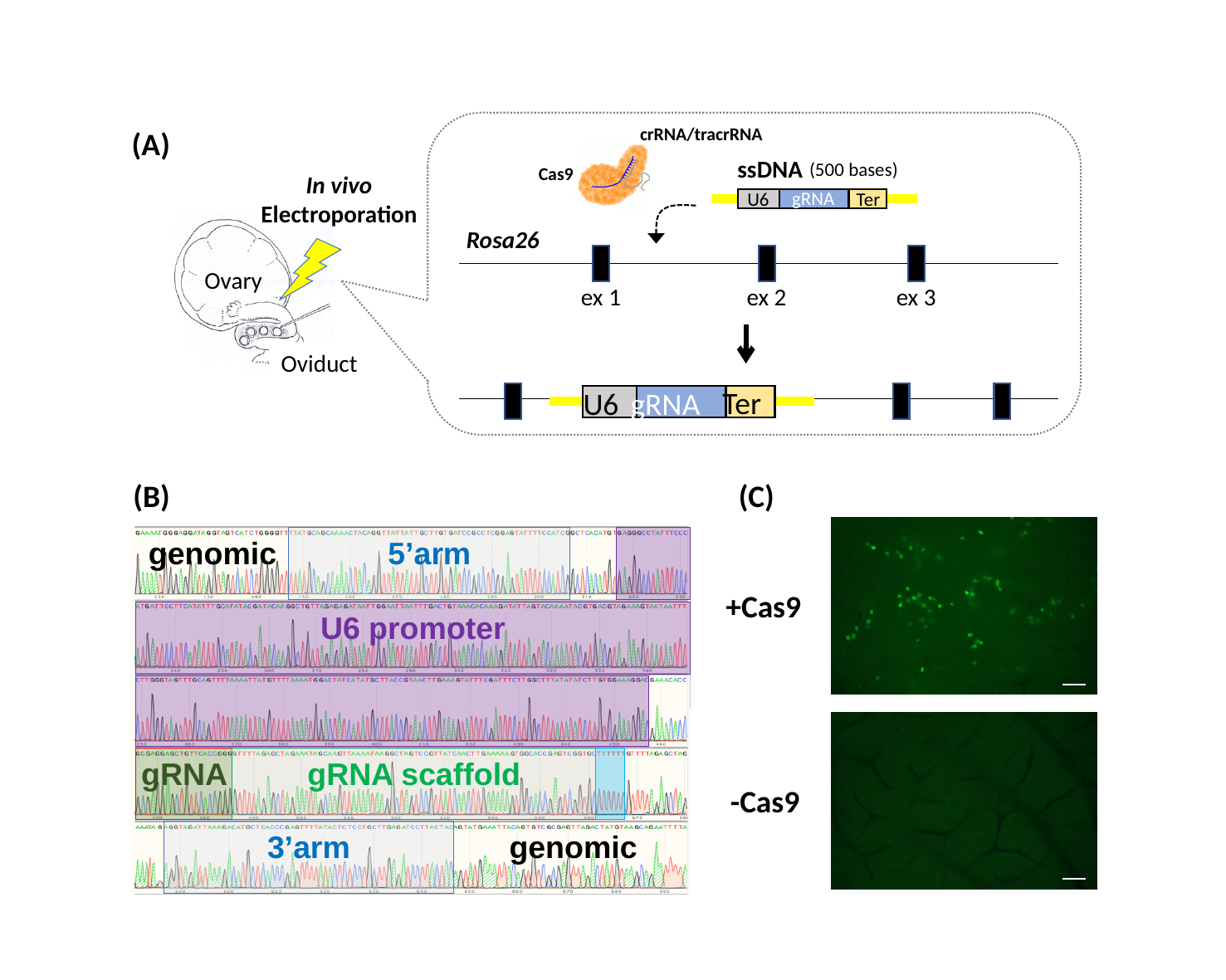

crRNA/tracrRNA
(A)
ssDNA
(500 bases)
Cas9
In vivo
Electroporation
Ovary
Oviduct
gRNA
U6
Ter
Rosa26
ex 1
ex 2
ex 3
Ter
U6
gRNA
(B)
(C)
genomic
5’arm
U6 promoter
gRNA
gRNA scaffold
3’arm
genomic
+Cas9
-Cas9
