## Supplementary material for "Delivering mRNAs to mouse tissues using the SEND system": Fig. S3

### Slide 1
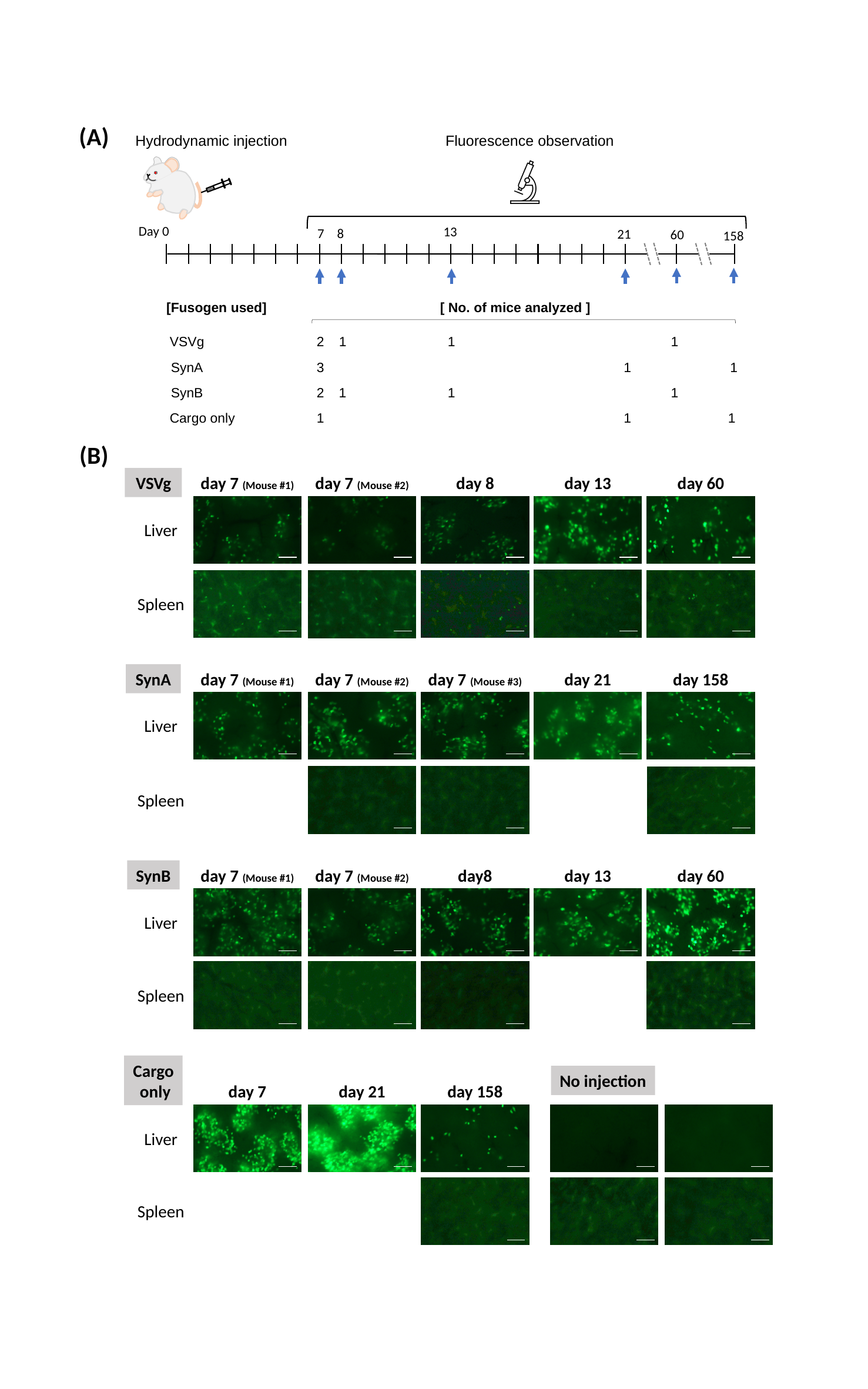

(A)
Hydrodynamic injection
Fluorescence observation
Day 0
13
7
8
21
60
158
[Fusogen used]
[ No. of mice analyzed ]
2
1
1
1
VSVg
3
1
1
SynA
SynB
2
1
1
1
1
1
1
Cargo only
(B)
VSVg
day 7 (Mouse #1)
day 7 (Mouse #2)
day 8
day 13
day 60
Liver
Spleen
day 7 (Mouse #2)
day 7 (Mouse #3)
day 21
day 158
SynA
day 7 (Mouse #1)
Liver
Spleen
SynB
day 7 (Mouse #1)
day 7 (Mouse #2)
day8
day 13
day 60
Liver
Spleen
Cargo
 only
No injection
day 7
day 21
day 158
Liver
Spleen
