## Supplementary material for "Delivering mRNAs to mouse tissues using the SEND system": Fig. S4

```
               .........10........20........30........40........50........60........70........80........90.......100
addgene        MAAAGGSSNCPPPPPPPPPNNNNNNNNTPKSPGVPDAEDDDERRHDELPEDINNFDEDMNRQFENMNLLDQVELLAQSYSLLDHLDDFDDDDEDDDFDPEP   100
NP_570947.2    MAAAGGSSNCPPPPPPPPPNNNNNNNNTPKSPGVPDAEDDDERRHDELPEDINNFDEDMNRQFENMNLLDQVELLAQSYSLLDHLDDFDDDDEDDDFDPEP   100

               .......110.......120.......130.......140.......150.......160.......170.......180.......190.......200
addgene        DQDELPEYSDDDDLELQGAAAAPIPNFFSDDDCLEDLPEKFDGNPDMLGPFMYQCQLFMEKSTRDFSVDRIRVCFVTSMLIGRAARWATAKLQRCTYLMH   200
NP_570947.2    DQDELPEYSDDDDLELQGAAAAPIPNFFSDDDCLEDLPEKFDGNPDMLGPFMYQCQLFMEKSTRDFSVDRIRVCFVTSMLIGRAARWATAKLQRCTYLMH   200
                                                                                    Retrotransposon gag protein (PF03732)

               .......210.......220.......230.......240.......250.......260.......270.......280.......290.......300
addgene        NYTAFMMELKHVFEDPQRREAAKRKIRRLRQGPGPVVDYSNAFQMIAQDLDWTEPALMDQFQEGLNPDIRAELSRQEAPKTLAALITACIHIERRLARDA   300
NP_570947.2    NYTAFMMELKHVFEDPQRREAAKRKIRRLRQGPGPVVDYSNAFQMIAQDLDWTEPALMDQFQEGLNPDIRAELSRQEAPKTLAALITACIHIERRLARDA   300

               .......310.......320.......330.......340.......350.......360.......370.......380.......390.......400
addgene        AAKPDPSPRALVMPPNSQTDPTEPVGGARMRLSKEEKERRRKMNLCLYCGNGGHFADTCPAKASKNSPPGKLPGPAVGGPSATGPERIRSPPSEASTQHL   400
NP_570947.2    AAKPDPSPRALVMPPNSQTDPTEPVGGARMRLSKEEKERRRKMNLCLYCGNGGHFADTCPAKASKNSPPGKLPGPAVGGPSATGPERIRSPPSEASTQHL   400

               .......410.......420.......430.......440.......450.......460.......470.......480.......490.......500
addgene        QVMLQIHMPGRPTLFVRAMIDSGASGNFIDQDFVIQNAIPLRIKDWPVMVEAIDGHPIASGPIILETHHLIVDLGDHREILSFDVTQSPFFPIVLGIRWL   500
NP_570947.2    QVMLQIHMPGRPTLFVRAMIDSGASGNFIDQDFVIQNAIPLRIKDWPVMVEAIDGHPIASGPIILETHHLIVDLGDHREILSFDVTQSPFFPIVLGIRWL   500

               .......510.......520.......530.......540.......550.......560.......570.......580.......590.......600
addgene        STHDPHITWSTRSIVFNSDYCRLRCRMFAQIPSNLLFTVPQPNLHPYLLHHVHPHVHPHMHQHLHQHLHQFLHPDPHQYPHPDPHYHHHQQADMQHQLQQ   600
NP_570947.2    STHDPHITWSTRSIVFNSDYCRLRCRMFAQIPSNLLFTVPQPNLHPYLLHHVHPHVHPHMHQHLHQHLHQFLHPDPHQYPHPDPHYHHHQQADMQHQLQQ   600

                                                                                          Deletion #1  Deletion #2
               .......610.......620.......630.......640.......650.......660.......670.......680.......690.......700
addgene        YLYQYLYYHLYPVMHHHLPPDQHEHLHEYLHQYLHQYLHQFLHHHLHPDLHQYLYQYLHNHMNPDPHHHPHPDPPQDPHHPPHQ------DPHQDPP---   691
NP_570947.2    YLYQYLYYHLYPVMHHHLPPDQHEHLHEYLHQYLHQYLHQFLHHHLHPDLHQYLYQYLHNHMNPDPHHHPHPDPPQDPHHPPHQDPHQHPDPHQDPPHQD   700
                                                                   consensus disorder prediction

               .......710.......720.......730.......740.......750.......760.......770.......780.......790.......800
addgene        ---------------------------------------------HQDPHQDAHQDPHMDPHLHQHQHPQPQPHPQQHPNHPQQPPFFYHMAGFRIYHPV   746
NP_570947.2    PHQHPDPHQDPPHQDPHQHPDPHQDPPHQDPHQDAHQHQDPHQDAHQDPHQDAHQDPHMDPHLHQHQHPQPQPHPQQHPNHPQQPPFFYHMAGFRIYHPV   800

               .......810.......820.......830.......840.......850.......860.......870.......880.......890.......900
addgene        RYYYIQNVYTPVDEHVYPGHRVVDPNIEMIPGAHSLPSGHLYSMSESEMNALRNFVDRNVKDGLMTPTVAPNGAQVLQVKRGWKLQVTYNCRAPQSGTIQ   846
NP_570947.2    RYYYIQNVYTPVDEHVYPGHRVVDPNIEMIPGAHSLPSGHLYSMSESEMNALRNFVDRNVKDGLMTPTVAPNGAQVLQVKRGWKLQVTYNCRAPQSGTIQ   900

               .......910.......920.......930.......940.......950.......960.......970.......980.......990......1000
addgene        NQYLRMSLPNMGDPAHLASYGEFVQVPGYPYPAYVYYTSPHMMTAWYPVGRDVHGRIIVVPVVITWSQNTNRQPPVPQYPPPQPPPPPPPPPPPPPPPPA   946
NP_570947.2    NQYLRMSLPNMGDPAHLASYGEFVQVPGYPYPAYVYYTSPHMMTAWYPVGRDVHGRIIVVPVVITWSQNTNRQPPVPQYPPPQPPPPPPPPPPPPPPPPA  1000

                     HA tag
               ......1010.....
addgene        SSCSAAYPYDVPDYA   961
NP_570947.2    SSCSAA---------  1006
```
